# Methanotroph Dynamics at Landfill Cover Soil Methane Emission Hotspots

**DOI:** 10.1101/2025.08.27.672573

**Authors:** Emmanuelle Roy, Sandani A. Buddhima, Henry Gibbons, Maria Strack, Laura A. Hug

## Abstract

Landfills contribute significant emissions to the global methane cycle, where emission hotspots can account for the majority of methane released. Methanotrophs in landfill cover soils can mitigate these methane emissions, but are constrained by geochemical conditions in the soils and the climate at the landfill’s location. We sampled cover soils from four Ontario landfills with differing characteristics, including measurements of methane flux, soil methane concentration, and a suite of geochemical variables. Sampling sites were distinguished based on the levels of methane flux. Microbial community diversity and methanotroph dynamics were examined using 16S rRNA gene amplicon sequencing. Sites with high methane emissions showed strong enrichment of methanotrophs, dominated by the genus *Methylomicrobium*. The distribution of methanotrophs in samples across landfills and sites differed significantly when considering community evenness. Of the environmental factors examined, microbial community diversity correlated most strongly to nitrate and nitrite concentrations. Methanotroph dynamics across different methane exposures and geochemical conditions within landfill cover soils inform the use of designer cover soils and/or methanotroph amendments in efforts toward mitigating methane emissions from landfills.

## Introduction

Methane (CH_4_), a potent greenhouse gas in the atmosphere, contributes to climate change (1). In Canada, landfills produced 19% of global anthropogenic methane emissions in 2023 (2), with methane emission from landfills predicted to increase alongside increasing waste generation globally (3). Hotspots within landfill cover soils are regions where abnormally high levels of methane emission are observed. Hotspot sites have been shown to contribute up to 73% of the total methane released at a landfill, even when comprising only 10% of sampling locations, making them priority targets for methane emission mitigation (4).

Landfills are heterogenous environments hosting an abundance of diverse microbes (5). These microbial communities play important roles in the ecosystem, interacting with the deposited waste or the products from microbial metabolism of this waste (6). Net methane emissions from landfills are a result of the balance between methanogens producing methane in anoxic locations within the landfill and methanotrophs oxidizing this methane in oxic landfill layers (7,8). Methanotrophs in landfills are primarily found in cover soils, where there is both sufficient oxygen and a supply of methane (9).

Within the methanotroph guild, genera are categorized based on carbon assimilation pathway and methane affinity. Type I methanotrophs use the ribulose monophosphate pathway for carbon assimilation (10). Type I methanotrophs are typically found in regions with lower methane flux and within an alkaline pH range (11). In contrast, type II methanotrophs use the serine pathway for carbon assimilation and are found in environments with high methane fluxes and at more acidic pH (11). Anaerobic methanotrophic archaea (ANME) consume methane under anoxic conditions using a reverse methanogenesis pathway (12). The oxidation of methane in non-ANME methanotrophs is mediated by the enzyme methane monooxygenase (MMO), which has two forms. Particulate methane monooxygenase (pMMO) is present in almost all methanotroph species, and is anchored within the intra-cytoplasmic membrane of the cell (11,13). In contrast, soluble methane monooxygenase (sMMO), which is localized to the cytoplasm of the cell, is observed in comparatively few methanotroph species. Both MMO forms allow methanotrophs to oxidize methane (11,13).

The geochemistry and physical properties of a given soil drive microbial community distribution and impact methane oxidation by methanotrophs (14). Soil pH is an important environmental factor for methanotrophs, where pH above 7.6 allows for optimal growth and methane oxidation for type I methanotrophs, whereas type II methanotrophs oxidize methane optimally at pH less than 7.0 (15). Optimal methane oxidation is observed when temperatures are between 30-40 degrees Celsius (16) and when gravimetric soil moisture content is between 20% to 35% (17). The presence of ammonium can be an inhibitor to methanotrophs as it competes with methane in the MMO enzyme, where oxidation of ammonium creates nitrite and toxic hydroxylamine (18). A similar inhibitory effect occurs with specific anions and cations, such as chlorides, nitrates, and bromides (19). Other components in the soil, such as the total organic carbon and total nitrogen, as well as the total inorganic carbon can mediate oxidation rates for methane (20,21). Most landfill cover soils are selected based on availability and economy – changes in cover soil composition that better support key methanotroph lineages could have significant impacts on emissions.

Harnessing the specific methane oxidizing capabilities of methanotrophs could be an efficient method to reduce methane emissions from landfills (22), including identifying methanotrophs best suited to these environments. As an example, *Methylotuvimicrobium buryatense* 5GB1C is a methanotroph able to efficiently oxidize methane at methane concentrations of 500 ppm and lower, compared to most methanotrophs that require much higher concentrations (5,000+ ppm) (22). Methanotrophs have potential as targeted tools for the mitigation of methane emissions, but this approach has not yet been broadly applied (23).

Here, we determine the trends in total microbial community composition, the methanotroph community composition, and the geochemical conditions for hotspot locations and control sites (low emission areas) in landfill cover soils across four Ontario landfills over a five-month time span. We identified correlations between environmental factors, and which variables were most strongly associated with differences in community composition. We predicted methanotrophic potential across sites and assessed the phylogenetic diversity across soils exposed to different methane flux magnitudes. Our goal was to determine conditions and specific lineages of methanotrophs best suited to address hotspot locations.

## Materials and Methods

### Landfill Sampling

Four landfills were selected, all managed by and located within the Regional Municipality of Peel in Ontario, Canada. The first landfill, North Sheridan Landfill (SH), is a closed landfill located in the York region (Part of Lot 14/Credit Indian Reserve (site A: 43.531517,-79.647679, site B: 43.531422,-79.647760, site C 43.531387,-79.647523)). Britannia Landfill (BB) was converted into a golf course in 2002, and is now called BraeBen golf course, in Mississauga, Ontario (site A: 43.598169,-79.698331, site C 43.598257,-79.698377). The Chinguacousy Landfill (CH) is a closed landfill, located in the town of Caledon (Mississauga City, Regional Municipality of Peel, Ontario Lot 28, Concession 3 (site A: 43.803281,- 79.848517, site B: 43.803240,-79.848641, site C 43.803231,-79.848549)). The fourth landfill, Caledon Landfill (CA), is still active with several closed and capped cells, which is located in the town of Caledon (1795 Quarry Drive, Lot 15, Concession 3 (site A: 43.835579,-80.015834, site B: 43.835527,-80.015802, site C 43.835661,-80.015772)).

At each landfill location, sites on capped cells with cover soil and established plant cover were chosen based on the observed methane emission levels. One or two hotspot sites were identified where local methane concentrations were at least threefold greater than atmospheric methane concentrations. Hotspot sites were labelled XXA and XXB (e.g., CAA for the Caledon landfill site A; note the BB landfill does not have a BBB site as only one hotspot was identified). Alongside hotspot sites, control sites (XXC) were chosen based on no measured methane emission or reduction of methane concentrations during chamber measurements (i.e., oxidation of atmospheric methane). Control sites were selected to be at similar elevations and with similar plant cover to hot spots, often within 10 m of the identified hotspots.

Sample collection and measurements were performed along a transect at each site, with four points along the transect chosen at 0 m, 0.5 m, 1 m, and 2 m from either the highest point of methane efflux (hotspot) or an arbitrarily chosen start point (controls). This resulted in a total of 44 samples across all four landfills for each sampling month. Sampling was conducted starting in late June 2023, and continuing in the last half of the month for July, August, and October 2023. A total of 176 samples were taken across all landfills in total.

Soil samples were collected at each of the four points on a transect, with approximately 100 g of cover soil (excluding any vegetation and large rocks) collected using a trowel, sterilized using 70% ethanol between each use to avoid cross contamination. The soil was placed in a sterile 24 oz whirl pack bag, sealed, and stored within a large cooler containing icepacks until soil samples could be stored in a fridge at 4 °C before processing. Soil was processed within 2-3 days of collection.

### Methane Flux Measurement and Soil Gas Collection

Methane flux measurements at each point on the transect were performed using the LI- 7810 CH_4_/CO_2_/H_2_O Trace Gas Analyzer (LI-COR). Four circular plastic collars (20 cm in diameter) were placed along the transect at each point (0 m, 0.5 m, 1 m, and 2 m). These collars were hammered several centimeters into the soil with a mallet to secure them firmly within the soil to prevent any gas leakage. To begin measurements, a closed acrylic chamber (20.5 cm diameter and 39.5 cm high) outfitted with an interior fan to circulate air was placed on top of the collar. Chamber temperature was measured with a type-K thermocouple. The chamber had inlet and outlet connections to the LI-COR analyzer, where the change in methane concentration (in ppb) in the chamber was recorded. The chamber remained on the collar for the duration of the measurement, until the methane concentration had reached an asymptote indicating saturation of flux, usually about three minutes for control sites and six to ten minutes for hotspot sites. The methane flux was calculated from the linear slope of the change in methane concentration over the closure, determined using the PEDRO software (https://github.com/BrianNewton/PEDRO/blob/master/CITATION.cff) and correcting for chamber volume and temperature.

Soil gas was retrieved using soil probes and 20 mL gas tight syringes. Hollow aluminum tubes were inserted into the soil to 10 cm depth next to the chamber at each point on the transect. Tubes were capped with an airtight sampling port, and flushed by removal of 30 mL of gas three times, ensuring dead volume was removed and soil pore gas was sampled. A volume of 15-20 mL of gas was extracted from the soil and inserted into 12 mL pre-vacuumed exetainers. Gas chromatography/mass spectrometry (GC/MS) was performed to determine the concentration of methane (ppm) for each sample.

### Soil Physical Properties and Soil Chemistry

The gravimetric soil moisture and pH were measured at the 0 m transect location at each site, with values presumed to be consistent across the transect. For soil moisture, 10 g of soil was weighed and dried in an oven at approximately 45 °C for 36 to 48 hours. The gravimetric soil moisture was calculated using the weight of the dried soil and its original weight, following O’Kelly and colleagues (24). The pH was determined using 5 g of the same dried soil sample mixed with 45 mL of MilliQ water using a pH probe, following Awotoye and colleagues (Awotoye et al., 2011). The temperature of the soil was determined using a temperature probe, which was inserted to 10 cm depth next to the chamber at each point along a transect and allowed to equilibrate.

Pore water extractions were conducted on 4.5 g of soil mixed with 45 mL of MilliQ water. After mixing, the sample was separated using the ThermoXTR centrifuge for 10 minutes at 4 °C, from which the supernatant was collected and filtered through a 0.45 𝜇m PES membrane for analysis. The extraction procedures followed that of Zhang and colleagues, with changes to the type of water used, the soil to water ratio and the time and temperature settings of the centrifuge (25). The analysis of filtered samples was performed by the Ecohydrology lab at UWaterloo. The collection of filtered pore water samples were kept and analyzed in acid washed vials or glassware. The inorganic carbon (IC) and the dissolved organic carbon was quantified by collecting 7 mL of the filtered extracted water, with the TOC-L IC reactor kit, which allows for sparging of the sample and the Nondispersive infrared detector quantifies the inorganic carbon from the carbon dioxide that it detects. This was kept at 4 °C until analysis. The dissolved organic carbon and total nitrogen (DOC/TN) was quantified using 7 mL of the filtered extracted water and the addition of three drops of hydrochloric acid (1 M) and the pH was taken to confirm it was within the range of two to three, storing this at 4 °C until analysis. Analysis of both IC and DOC/TN were performed in accordance with the Shimadzu Total Organic Carbon Analyzer TOC-L CPN User’s Manual (Shimadzu, Canada). To quantify cations and trace metals present in the sample, a 10 mL subset of the filtered extracted water was mixed with 2% ultrapure nitric acid and kept at 4 °C until analysis. The quantification was carried out using a Thermo Scientific iCAP 6300 Duo ICP and the CETAC Technologies ASX-520 autosampler by Inductively Coupled Argon Plasa Optical Emissions Spectrometry (ICP OES) (Thermo Fisher Scientific, Canada). The anions present in the sample were determined using ion chromatography, employing the Dionex ICS-5000 Capillary Ion Chromatography system (Thermo Fisher Scientific, Canada). 1 mL of the filtered extracted pore water was further filter using a 0.2 𝜇m polypropylene filter and stored at -20°C until analysis, following the procedures outlined in the manual for the IonPac AS-18 Dionex system (Thermo Fisher Scientific, Canada). Ammonium levels (NH ^+^) were found using a Gallery Discrete Analyzer from a 5 mL sample of the filtered extracted pore water, measuring the absorbance correlating to the concentration of ammonia present (Thermo Fisher Scientific, Canada). Analysis of cations, anions, and ammonium levels were performed in accordance with the Thermo Fisher Scientific User’s Manual (Thermo Fisher Scientific, Canada).

### DNA Extraction and Amplicon Sequencing

Microbial genomic DNA was extracted from the soil samples using the DNeasy PowerSoil Pro kit (QIAGEN, Canada) following the manufacturer’s protocol with no modifications. Aliquots of extracted DNA were sent to MetagenomBio for 16S rRNA gene amplicon sequencing on a MiSeq sequencer, using paired end Illumina sequencing and 2 x 250 reads. Forward and reverse primers targeted the V4 region of the 16S rRNA gene; 515F (5’ GTGYCAGCMGCCGCGGTAA 3’) and 806R (5’ GGACTACNVGGGTWTCTAAT 3’) (26). To test for contamination, kit controls excluding the soil sample were extracted and sequenced alongside the true samples. Sequencing controls excluding template DNA were also included following the same sequencing procedure.

### Sequence Analyses

Initial amplicon sequencing sample processing was conducted in QIIME2 (27). The raw sequence files from MetagenomBio were processed using the QIIME2 software package to generate microbial community representations for each sample, for taxonomic identification, and for visualization (27). The reads were trimmed 35 base pairs from each end of the sequence to remove the primers and low-quality sequence at read ends (27). Amplicon sequence variants (ASVs) were determined using DADA2 (28). A trained SILVA database (version 138) was used to identify the amplicon sequence variants (ASV)s for taxonomic assignment (29). All statistical analyses, manipulation, and visualization of the sequence data were conducted using R statistical software v4.3.2 (30). The bar plots for methanotroph populations were made using the taxonomic identification performed by QIIME2, exported to R, and visualized using *ggplot*. The methanotroph correlation plot was made using *ggplot2* and *ggmisc* (31). All PCoAs, Shannon Index, and observed scatter plots were constructed and visualized using *phyloseq*, *ggplot2*, and *qiime2R* (31–33). The PCoA plots with overlaid arrows was generated using the R packages *vegan*, *ggplot2*, *qiime2R*, *dplyr*, and *data*.*table* (31,33–36). Changes in geochemical data across sites and landfills were visualized using R studio, using the R package *ggplot2* for scatter plots and boxplots (31). The Kruskal-Wallis rank sum test was used to test for statistical significance between measurements as values were not normally distributed (37). Spearman correlation analysis was calculated using the ggpubr package in R to investigate the relationships between the geochemical data (38), using p < 0.05 to indicate significance (39). A Benjamini-Hochberg correction was applied to p-values to account for multiple tests (40).

### PICRUSt2 Analysis

To assess the potential proportions of the methane monooxygenase enzymes in the microbial communities present, the PICRUSt2 plugin in QIIME2 was used with ggPICRUSt2 . The PICRUSt2 pipeline was followed using the raw reads of the sequences, and visualizations were done using the R studio packages *qiime2R*, *phyloseq*, *dplyr* and *ggplot2* (31–33,35).

## Results

### Geochemical Conditions

The methane flux was significantly higher for sites identified as hotspots compared to control sites (p = 2.2x10^-16^), with an average flux value of 87,024.44 mg CH_4_ m^-2^ d^-1^ for hotspot sites (ranging between -19.11 mg CH_4_ m^-2^ d^-1^ and 1,167,261.02 mg CH_4_ m^-2^ d^-1^) and an average flux value of 109.28 mg CH_4_ m^-2^ d^-1^ for control sites (ranging between -22.30 mg CH_4_ m^-2^ d^-1^ and 2185.19 mg CH_4_ m^-2^ d^-1^). This was expected given the site categorization parameter was methane emissions, but which remained stable over the timeframe of this survey. The methane flux also varied along the 2 m transects, with some points along a given transect having large differences in methane flux compared to other points in the same transect. As an example, for CAA in August, the methane flux at the initial site of the hotspot (0 m) was very high (798,966.23 mg CH_4_ m^-2^ d^-1^), whereas at the 2 m mark the methane flux was down to 71,823.61 mg CH_4_ m^-2^ d^-1^ (Figure 1). Methane flux values were generally higher closer to the 0 m mark for each transect with a hotspot, but not universally (Figure 1).

**Figure 1.**
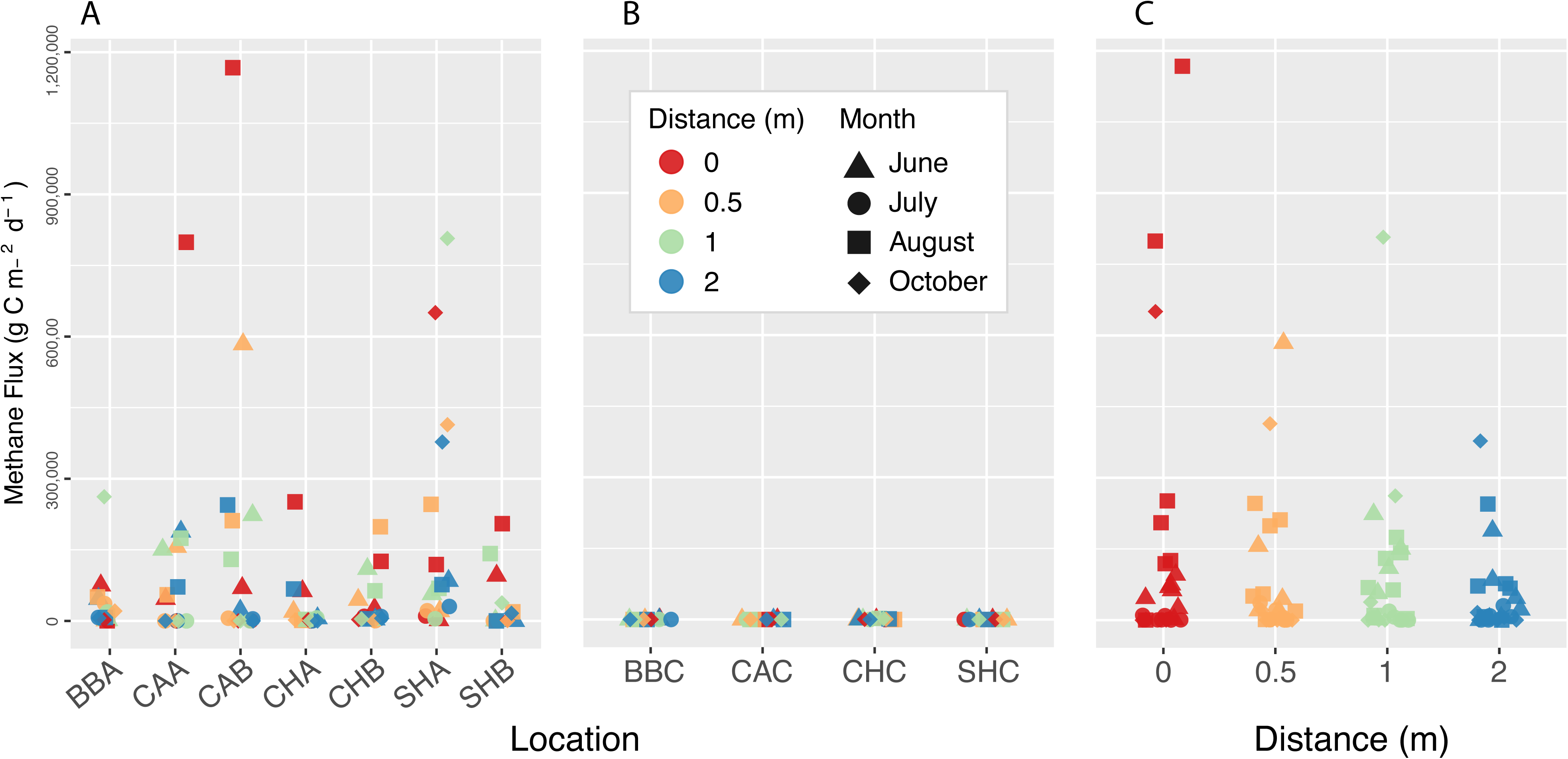
Methane flux values determined from *in-situ* measurements using the LiCOR gas analyzer and a known-volume chamber, with plots A and B depicting differences across hotspot and control sites, respectively, and plot C focusing on the change in flux along hotspot transects. Samples are separated by site in A and B, with month of sampling denoted by shape and transect distances by colour (N=176). The Kruskal-Wallis rank sum test identified significant differences between locations and sample type (hotspot vs. control), with p values of 2.53x10^-12^ and 1.16x10^-15^, respectively. Site names indicate the landfill and site type (hotspot sites XXA,XXB; control site XXC).

The geochemical conditions (*i.e.,* pH, soil moisture, soil methane concentration, soil temperature) were measured once per transect for each condition across all months, while soil chemistry (*i.e.,* dissolved organic carbon (DOC), dissolved inorganic carbon (DIC), total nitrogen (TN), Ammonium, Nitrite, Nitrate, Aluminum, Calcium, Chloride, Fluoride, Iron, Magnesium, Manganese, Phosphate, Phosphorus, Potassium, Silicon, Sodium, Sulfur, Sulphate) was determined for the 0 m transect location for all sites for the months of August and October. The gravimetric soil moisture varied more for the hotspot sites compared to the control sites. The moisture content ranged between 5.7–118.6% with a median of 22.4% for hotspots, compared to a range of 9.2–36.1% and a median of 20.8% for control sites (Figure S1). The Kruskal-Wallis rank sum test depicts a significant difference across site locations (e.g., XXA, XXB, XXC), with a p value of 0.048. Soil moisture was not significantly different between control and hotspot sites (p=0.479, Figure S1). The pH across each site for all four months varied across landfills and site locations, with significant differences between site landfills (p = 0.001) and site types (p=9.43x10^-5^, Figure S2). Notably, pH ranges for the hotspot sites were closer to a neutral value of 7.0 (range 6.4–7.9, median 7.3) while control sites had a more alkaline pH range (range 7.5– 8.1, median 7.8; Figure S2). Soil temperature ranged from 18.1°C to 33.2°C; hotspots had on-average significantly higher temperature (hotspot mean = 23.9°C, control mean = 20.3°C; Kruskal-Wallis, p = 2.11x10^-5^) (Figure S3).

To characterize variation in soil chemistry we examined a suite of cations and anions, and measured total nitrogen as well as dissolved organic carbon (DOC) and dissolved inorganic carbon (DIC). There were no clear differences between control and hotspot sites for any variable, though site CHA had distinctly higher concentrations for DOC, total nitrogen, chloride, calcium, magnesium, manganese, and sulfur, with all but chloride more strongly elevated in the October sample compared to August (Figure S4). Other sites showed sporadic elevation of one or a few variables, with no consistency between months for these compounds (Figure S4). DOC concentration was relatively low across all samples (mean = 0.412 mg C/g soil (dry), range 0.073–3.805 mg C/g soil (dry)), excluding CHA, while DIC was uniformly low (mean = 0.070 mg C/g soil (dry), range 0.040–0.134 mg C/g soil (dry)) (Figure S4).

Spearman ranked correlation tests identified 21 variable pairs with significant correlations for the full dataset, following Benjamini-Hochberg adjustments of p-values for multiple tests (p<0.05) (see full test results in Supplementary data file 1). For the hotspot only data, 25 pairs of measured variables were significantly correlated. Only one variable pair (Aluminum and Iron) was significantly correlated for the control dataset (p = 0.011), where Aluminum and Iron were also significantly correlated for the full and hotspot-only datasets (Table S1). The low number of significant variables for control samples was likely due to too low a sample number. Of the significant pairs, 12 were consistent between the full and hotspot-only datasets (Table S1, Supplementary Data File 1). Notably, for the measured geochemical conditions, soil moisture was not correlated with any other variable, while methane flux and soil methane concentration were positively correlated for the full (spearman’s rho: 0.73) and hotspot-only datasets (rho: 0.55; note, control site methane flux values were almost exclusively 0, precluding statistical comparison) (Table S1, Supplementary Data File 1). Methane flux and soil temperature were significantly positively correlated in both the total and hotspot datasets (rho: 0.41, 0.33, respectively), and soil methane was positively correlated with soil temperature for the total dataset only (rho: 0.47). Methane flux and methanotroph proportional abundance were significantly correlated only for the total dataset (rho: 0.28, p=0.0069). Nitrate was significantly negatively correlated with methane flux in both the full (rho: -0.77) and hotspot (rho: -0.78) datasets (Table S1), and with soil gas for the full dataset (rho: -0.60). Hotspots showed significant negative correlations between methane flux and nitrite concentrations (rho: -0.78) and flux with sulphate (rho: -0.78). Nitrite and Sulphate were strongly positively correlated (rho: 0.88) in the hotspot dataset, making it difficult to determine the driving factor behind these shared correlations with methane flux. Soil chemistry variable pairs made up the other significant correlations, with all significant pairings positively correlated with each other (See Table S1 for details on all significant pairwise comparisons).

### Microbial Community

The dominant microorganisms across all samples were from the families *Beijerinckiaceae* and *Methylococcaceae*, in that order. *Beijerinckiaceae* was present at a mean relative abundance of 10.5% (range 0–51.7%), more specifically appearing in higher numbers in sites BBA, CAA, CAB, CHA, CHB, CHC, SHA, and SHB. *Methylococcaceae* was present across all sites except control sites CAC, CHC and SHC, at a mean relative abundance of 9.5% (range 0–46.7%).

The methanotrophic component of the community was determined across all transects, with *Methylomicrobium* and *Methylobacter* as the two most abundant genera for each landfill (Figure 2). The most abundant methanotroph identified was *Methylomicrobium*, observed in all hotspots, while control sites contained a stronger *Methylobacter* signal within a lower overall abundance of methanotrophs. The mean relative proportion of methanotrophs at hotspots was 12.5% (range 0.14–41.3%), and for control sites was 5.7% (range 0–29.7%) (Figure 2). The proportion and type of methanotrophs stayed relatively consistent across all four months of sampling within sites (Figure 2). The control site for the BB landfill (BBC) was disrupted in August 2023 during golf course maintenance, therefore the transect was moved to a new site ∼5 m uphill from its original location, which impacted the time series. The observed methanotroph abundance at the BBC site in August occurred despite no measured methane emissions at either control location (Figure 2).

**Figure 2.**
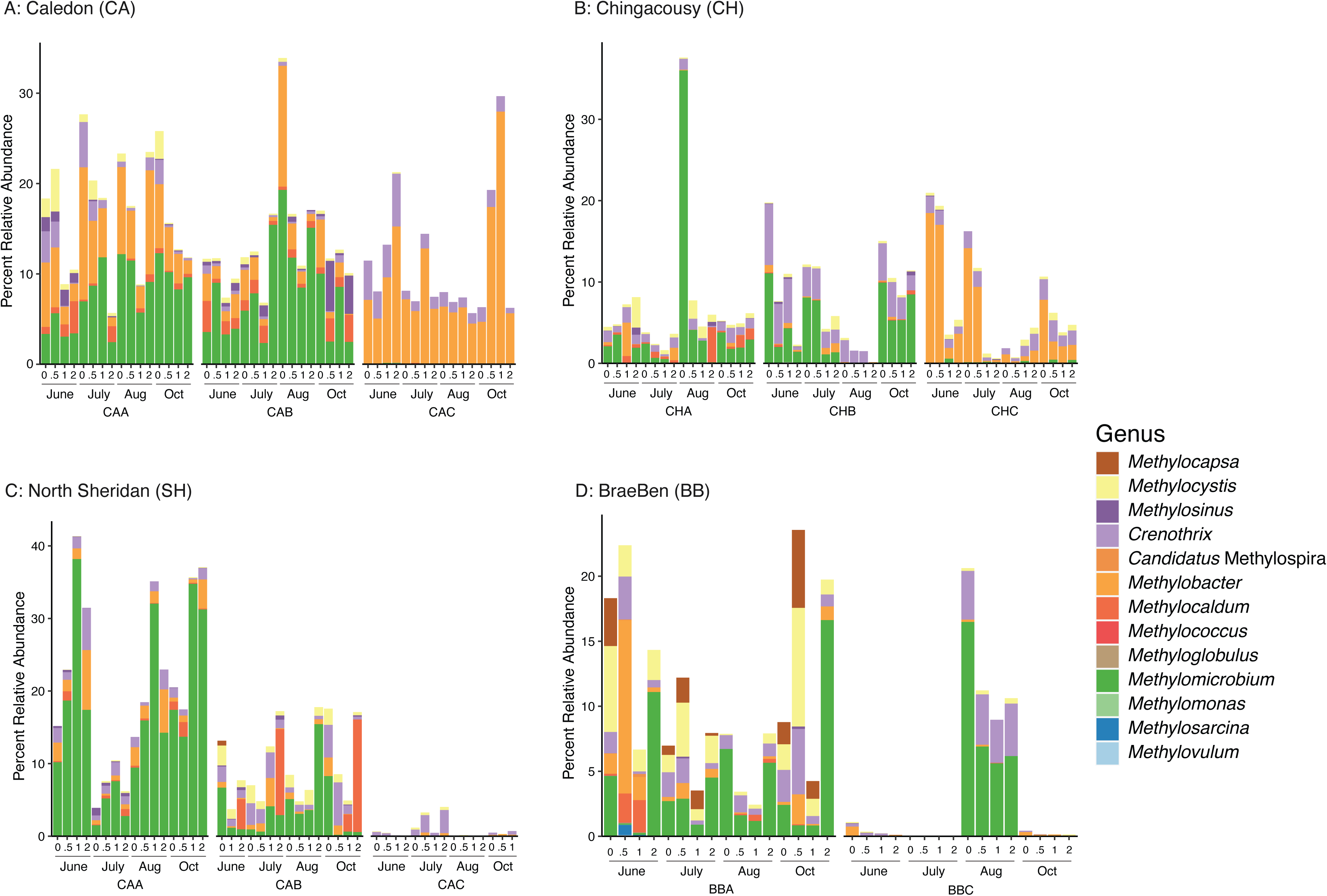
The relative abundance of methanotroph genera in each sample (N=174 samples). The four plots, A, B, C, and D, depict the four landfills, SH, CA, CH, and BB, respectively. Facets within each plot depict individual sites. Sample names indicate the Landfill, sampled site, date sampled, and transect distance from the hotspot (sites A,B) or control site (sites C) in meters. For example, “SHA.23.06.0.0” is the SH landfill, site A (hotspot), sampled June 2023, at the 0-meter mark.

The linear correlation between percent relative abundance of methanotrophs and methane flux value was assessed for the total methanotrophs dataset (Figure S5A) as well as for *Methylomicrobium* alone, as it was the genus present at the highest abundance (Figure S5B). Each landfill was found to have different R^2^ values corresponding to the correlation between proportion of methanotrophs and methane flux. No landfills were found to have a significant correlation between these two variables for either total methanotrophic community or for *Methylomicrobium* (max R^2^ = 0.15; Figure S5). This is in contrast to the overall Spearman’s rank correlation test, which identified a significant correlation between methanotroph proportion and methane flux for the total dataset (rho: 0.278, p=0.006).

The community alpha diversity was assessed with Observed and Shannon Index diversity metrics (Figure S6A). Kruskal-Wallis tests identified significant differences across sites (Shannon; p = 5.03x10^-16^, Observed; p = 0.002) and between control vs. hotspot (Shannon; p = 7.76x10^-19^, Observed; p = 4.04x10^-6^), but not between landfills (Table S2). The Kruskal-Wallis test was repeated for the methanotroph community specifically, identifying significant differences across sites and landfills for the observed diversity only (Site; p = 0.024, Landfill; p = 0.025) (Table S2, Figure S6B). The Shannon index values ranged from 2.7 to 6.9, with highest values seen in BBA at the 0 and 0.5 m mark on the June transect. For methanotrophs specifically, these values fell between 0.9 (almost none detected) and 2.9 (Figure S6). Most metrics showed methanotroph diversity decreasing over time, with lowest values for each transect location in October. Methane flux and methanotroph diversity (as Shannon Index values) showed no correlation (R^2^= 0.037).

Beta diversity was calculated using weighted and unweighted Unifrac diversity metrics, exploring differences in the microbial communities across sites and between landfills (Figure S7). Statistical tests such as multivariate permutational analysis of variance (PERMANOVA) and multivariate permutational analysis of dispersion (PERMDISP) were used to test for significant differences between site, site type, and landfill for both the total microbial community as well as the methanotrophic community (Table S2). A weighted Unifrac metric using the methanotroph community dataset was the only test showing a significant difference between samples. This contrasts with the unweighted Unifrac measure, which considered only the richness of the methanotroph communities, which showed no significant difference between samples across either sites or landfills (Table S2). A Bray-Curtis PCoA was used to visualize community beta diversity overlaid with geochemical data for the total community (Figure 3A) and for the methanotroph fraction (Figure 3B). The strongest associations were observed for nitrate and nitrite concentrations for both datasets.

**Figure 3.**
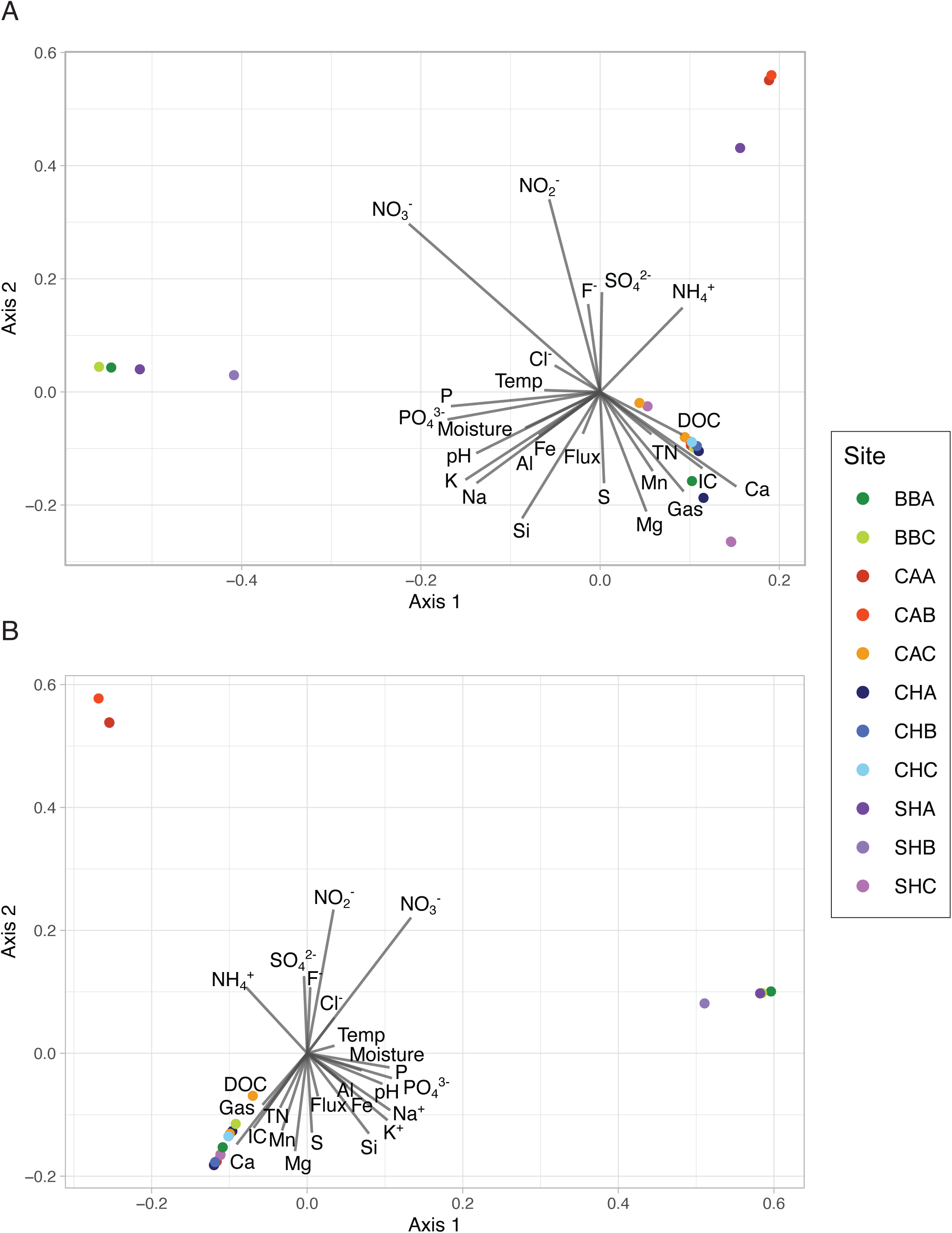
PCoA plots based on Bray-Curtis beta diversity measures for total (top) and methanotroph (bottom) diversity with geochemical variables overlaid, (N=174). Arrow length is proportional to the corresponding squared correlation coefficient value for a variable. The strength of an association between a geochemical condition and the change in microbial community composition is represented by the length of the arrow. Site names are as in Figure 1.

### Potential Function of Methanotrophs

The potential for different molecular mechanisms for methane oxidation was predicted using PICRUSt2 for the pMMO and sMMO enzymes. The capacity for methane oxidation was observed to be higher for hotspot sites compared to control sites for the pMMO enzyme (Figure 4). The sMMO enzyme was predicted in an average of 0.42% of methane oxidizing lineages across all sites, with the SHB site reporting 0 predicted sMMO-containing cells across all transects and throughout all four months (Figure 4). In comparison, the pMMO enzyme was predicted to be present, on average, in 26.3% of methane oxidizing lineages across sites, with each site, including all control sites, demonstrating predicted pMMO activity.

**Figure 4.**
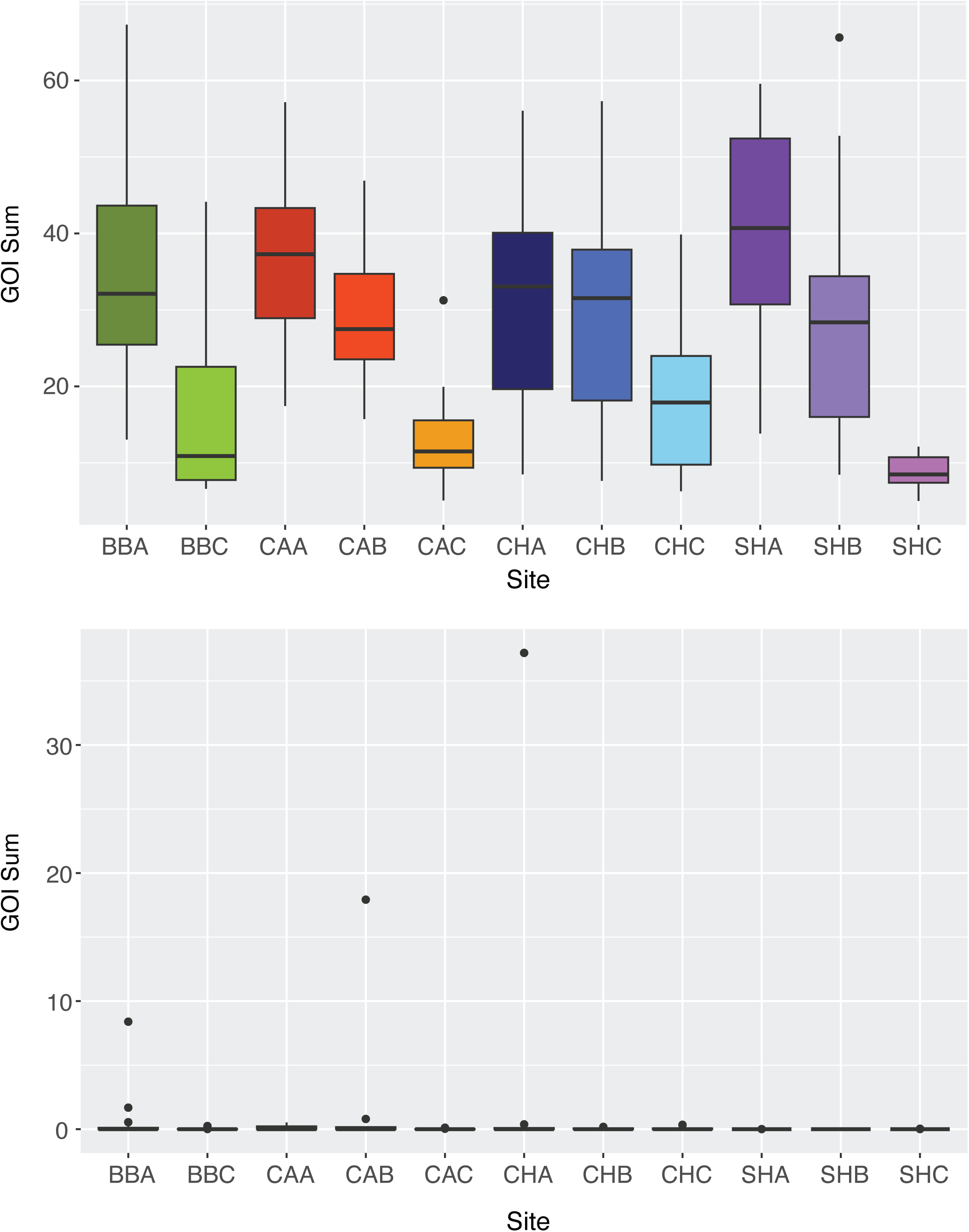
The predicted potential for methane oxidation performed by the methanotrophs present at each of the landfill sites, (N=174). The GOI Sum is the sum of ASVs per site derived from lineages predicted to encode the protein of interest, here particulate methane monooxygenase (top) and soluble methane monooxygenase (bottom). Box plot borders indicate the first and third quartile range of values, with the line indicating the mean. Outlier values are displayed as points. Site names are as in Figure 1.

## Discussion

### Variation in Geochemical Conditions

Geochemical conditions of landfill cover soils vary greatly, impacting the microbial diversity these environments support (14). To understand the driving factors that affect microbial diversity, we first examined the change in geochemical conditions across sites, comparing these factors between control and hotspot sites.

The methane flux changed significantly across site locations and site types, with flux values near zero for control sites and a large observed range for hotspot sites (Figure 1). Flux only occasionally followed a decreasing trend along a transect, as seen in the SHA site for the month of October (Figure 1). Transects that did not follow this expected pattern of reducing methane emissions with distance from the hotspot source likely contain multiple hotspot sites along the transect, especially in regions with fissures or cracks rather than point-source hotspots. The spearman’s ranked correlation was significant for the relative proportion of methanotrophs and methane flux for the full dataset only, but the linear correlation between these variables was very weak (R^2^= 0.05-0.12; Figure S5A, Table S1). The linear correlation between methane flux and the relative proportion of *Methylomicrobium,* the dominant methanotroph genus observed, was also weak (R^2^ = 0.06-0.15; Figure S5B). This observation does not agree with literature, as previous studies have shown a linear relationship between the proportion of methanotrophs and the concentration of methane or magnitude of methane flux in soils (41). Based on the extremely high flux values at the landfill hotspots, methane may represent an essentially unlimited resource, divorcing methanotroph abundance from resource availability. Complicating this is the fact that methanotrophs can persist in systems with limited or no methane for some time, and as such can be used as markers for legacy methane-emitting sites (42). Methanotroph persistence following the removal of a methane source is a parsimonious explanation for the methanotroph abundances at the CHC and CAC control sites, but their status as legacy hotspots is speculative.

Von Fischer & Hedin (2007) reported a positive correlation between elevated levels of soil moisture and increased net methane emission (43). When waste soil cover includes a high moisture content, the available pore space for gaseous transport and diffusion decreases (44), reducing the rate of methane oxidation due to constraints on methane and oxygen transport (45). Conversely, at low soil moisture contents, low oxidation rates are the result of a physiological reaction to water stress, which results in decreased microbial activity (46). Maximum methane oxidation rates are expected at a moderate volumetric moisture content of 15% (46). Both control sites and hotspot sites had median soil moisture content slightly above 15% (Hotspot = 22.4%, Control = 20.8%), which indicates that the methanotrophs present could be oxidizing methane at or near their maximum rate. Some hotspots sites experienced much higher soil moisture content (30%+), specifically the BBA site in June, and CAB, CHA, and CHB in July. Surprisingly, soil moisture was not correlated with any other measured variable, including methanotroph abundance. This lack of correlation is likely also due to methane saturation of the system given extremely high methane flux values. In this system, more saturated soils (30%+ moisture) still provide excess methane despite reduced diffusion rates, divorcing methanotroph abundance from soil moisture trends.

The soil pH differed significantly between site types and location (Figure S2), with control sites having a narrower and more alkaline range (Figure S2). Reddy and colleagues observed a shift in methane oxidation with varying pH values where the rate of methane oxidation was at its optimal value when the pH of the soil was 7.6 (15). This value is close to median pH for the two site types (Hotspot = 7.2, Control = 7.8). Reddy and colleagues also determined that type I methanotrophs dominated in environments with alkaline pH levels (between 7.6 and 10). The type II methanotrophs were abundant at pH levels below 7.0 (15). Most of the landfill sites are dominated by type I methanotrophs, despite the pH for most hotspot sites falling below 7.6 (19 below, 7 above, Figure S2). *Methylocystis* and *Methylosinus* are type II methanotrophs with higher abundances at BBA and CAA, sites with lower pH values across time (Figure 2, Figure S2). In contrast, CHA and SHB have low pH measurements for one month only, and do not show enrichment of type II methanotrophs. Soil pH is likely acting as a strong driving force dictating the genera observed (47,48), but consistent lower pH seems required to support a type II methanotroph competitive advantage in these soils.

Increased soil temperature promotes CH_4_ oxidation, up to a peak at 30 °C, with significant declines above 40 °C (49). Nearly all cover soil temperatures were below 30 °C in this study, with higher temperatures observed at hotspot sites compared to controls (Figure S3). Soil temperature did not significantly differ across landfills, but did between hotspot and control sites, with a general trend toward higher temperatures at hotspot sites, where the CH landfill was an exception to this trend (Figure S3). Methane flux and soil temperature were positively correlated in the total and hotspot datasets, with soil temperature being higher at hotspots likely driven by lower vegetation cover, as high methane concentrations damage plant roots and can lead to “dead spots” where the bare earth receives stronger solar radiation. The CH landfill hotspots were associated with trees, which reduced vegetation loss and provided shade, mediating soil temperatures. Soil temperature is known to impact methanotrophic communities, with strong shifts in community membership towards thermophilic lineages observed above 30 °C (50). Only hotspot sites at CA and SH experienced temperatures above 30 °C, in June, July and August, consistently showing high temperatures (>30 °C) at the 0 m point on the transect. Depending on the month, the high temperatures were sometimes consistent across the transect, or, in other instances, soil temperatures dropped by ∼10 °C across the two-meter transect. The methanotrophs identified from these higher-temperature soils are not from thermophilic lineages. This indicates methanotrophs at Ontario landfills have likely been selected for temperature resilience, but across a more temperate thermal range. This observation has concerning implications for methane oxidation rates in a warming climate, as the established methanotroph communities are not well-adapted to prolonged temperatures above 30 °C.

The chemical composition of the soil showed differences across sites, site types, and months within the expected range of soil conditions (Figure S4). CHA was notable for markedly higher concentrations of many of the measured soil chemistry characteristics, including total nitrogen, DOC, calcium, magnesium, sulfur, and manganese, specifically for the August timepoint. Christophersen and colleagues reported that oxidation rates increased with increased organic matter concentration (51). It was not possible to determine microbial methane oxidation rates from our survey, but the lack of correlation between methane flux or methanotroph proportional abundance with either DIC or DOC for all datasets indicates these are not strong driving factors in the community composition.

### Microbial Community

The landfill cover soils showed highest observed microbial diversity in June, with lower, more stable observed ASV counts in July and August, and a partial recovery of diversity in October (Figure S6). This shift may be due to heat stress on the soil microbes or changes in soil moisture over the summer months. Hotspots and control sites were significantly different across all alpha and beta diversity measures (n=8), excepting PERMDISP comparisons under the Bray-Curtis model (Table S2). Hotspots were experiencing significantly different soil pH and soil moistures, both factors that strongly impact microbial community memberships.

Methanotroph diversity varied significantly between hotspots and control sites, further distinguishing the overall microbial communities (Figure 2). *Methylomicrobium* was the most abundant methanotroph genus present across all samples, and the most abundant genus at hotspot sites. This genus’ members are type I methanotrophs (52) that are commonly detected in landfill cover soils, but were not the dominant methanotroph in these environments in prior surveys (11,53). *Methylomicrobium spp.* prefer low methane emission sites (11,53), making this genus’ abundance at hotspots with extremely high methane emissions unexpected. *Methylomicrobium alcaliphilum* has been a focus in metabolic engineering for its capacity for methane oxidation (54); identification of high-methane active *Methylomicrobium* lineages may inform efforts to develop biological tools for methane mitigation.

Control sites were expected to contain low to no methanotrophs, but this was only observed consistently at the SH landfill, with very low relative abundances of methanotrophs for all months at SHC. Unexpectedly, both CHC and CAC control sites displayed strong methanotroph signals, although with different community compositions compared to their associated hotspot sites (Figure 2). The presence of methanotrophs at sites where very low or no detected methane is being emitted could indicate these are legacy emission sites, colonized by methanotrophs capable of persisting in their environment in the absence of methane.

Alternatively, it is possible the methanotroph communities at these sites are oxidizing all of the methane transiting through these cover soils, and represent an active, but low-methane comparator to the hotspots. The type I methanotroph genus *Methylobacter* was the most abundant methanotroph at all control locations (with the exception of BBC in August, discussed below), in keeping with prior observations of this genus at low-methane locations (Figure 2) (11,50). *Methylobacter* species may have higher tolerance to changes in methane availability, or be capable of persisting in the absence of methane (55). Our results indicate *Methylobacter* is not a useful marker taxon for identifying methane-impacted soils, as its abundances did not track with methane signals. The BB landfill control site (BBC) experienced a significant shift from very low methanotroph abundances in June and July to a much higher relative proportion of methanotrophs in August (Figure 2). The original BBC site was disrupted by golf course maintenance prior to the August sampling, and the BBC site was relocated ∼5 m uphill from the original site. The second site had shifted back to a low-abundance, *Methylobacter*-dominated methanotroph community in October (Figure 2).

The methanotroph observed diversity was more stable than the total microbial community across both time and sampling locations, with the majority of sites containing 1-16 methanotroph ASVs (4 exceptions out of 178 samples, Figure S6). Systems dominated by type I methanotrophs typically show decreased evenness (56). In our survey, the Shannon index for the methanotroph-only community ranged from 0 to 2.94, with no significant differences across landfills, sites, or site types (mean 1.34, Figure S6, Table S3). The low Shannon index values for the Ontario landfill cover soils’ methanotroph communities may be due to type I methanotroph dominance. When comparing beta diversity measures, only weighted unifrac measures showed consistent significant differences between sites (e.g., XXA, XXB) and between landfills (Table S3). No distinguishing signal was observed between hotspots and control sites, nor were there obvious landfill or site-related clustering on a PCoA (Figure S7). This contrasts the significant differences between total microbial communities across all metrics (landfills, sites, site types) for both weighted and unweighted Unifrac metrics, indicating the methanotroph fractions are more similar to one another and more stable than the overall landfill cover soil microbial communities.

For both the total and methanotroph communities, the strongest factors influencing beta diversity were nitrate and nitrite concentrations (Figure 3). The availability of nitrate and nitrite are expected to act as driving factors for microbial community composition due to the importance of soil microbial nitrogen cycling. Methane flux is typically correlated to the presence of methanogens and methanotrophs and therefore should play an important role in the diversity of methanotroph-dominated communities (23); in our data methanotroph abundance was significantly correlated with flux for the total dataset, but not for hotspot-only or control-only data. Other environmental factors, notably ammonium, calcium, and magnesium, explained components of the distribution of methanotrophs and overall community dynamics (Figure 3).

Ammonium can have an inhibitory effect on MMO activity, where addition of ammonium has been shown to shift the distribution of methanotrophs towards type I methanotrophs (18,19). Both calcium and magnesium impact microbial attachment and are known to influence microbial community structure (57). No single variable, nor a small subset of variables is sufficient to explain the methanotroph distribution in the landfill cover soil communities, and none of the soil chemistry metrics were significantly correlated with methanotroph abundance.

### Functional Role of Methanotrophs

PICRUSt2-based predictions of the function within the surveyed microbial communities indicated a much higher abundance value for the pMMO enzyme for all sites across the four landfills (Figure 4) (58,59). sMMO had a median of zero summed ASVs predicted to contain this gene (Figure 4). Comparing control and hotspot sites, hotspot sites consistently had a higher number of ASVs predicted to encode pMMO, but all control sites had non-zero numbers of ASVs with potential pMMO genes. Methanotrophs of all types have been observed to express pMMO, where sMMO is more restricted, with sMMO expression reported from only one type I methanotroph, *Methylomonas methanica* (60), and a few type II methanotrophs (11). ASVs identified as *Methylomonas* were observed in the landfill cover soil data, however this population was at very low relative abundance (maximum 0.04%, in BBA). The more abundant methanotrophic genera in the microbial community were *Methylomicrobium*, *Methylobacter*, and *Crenothrix*, all type I methanotrophs (61,62). From the type I methanotrophs dominating the soil environment and the very low abundance of the predicted sMMO, we expect the sMMO activity to be minimal compared to pMMO.

## Conclusions

Landfill cover soil methane hotspots represent specific, high value targets for methane mitigation. Hotspot soils demonstrated dynamics counter to current expectations for methanotroph-dominated soils (41,45,46), with no correlation between soil moisture and methanotroph abundance, and no linear relationship between methane flux and methanotroph abundance. Given the extremely high rates of methane efflux from hotspots, it seems likely that methane is saturating the system, removing typical constraints on methane diffusion and availability.

The dominant methanotrophic genus at all hotspot sites was *Methylomicrobium.* In contrast, control sites with no observed methane efflux either had very low (<5%) methanotroph abundances, or contained primarily *Methylobacter*. Differentiation between hotspot and control sites based on methanotroph identity was stronger than using methanotroph presence/absence or abundance. *Methylobacter* was specifically identified as a poor marker for active methane efflux from landfill cover soil, as this lineage persists in legacy sites no longer experiencing high methane efflux.

The geochemical factors associated with strongest microbial community change were methane flux, nitrate and nitrite concentrations, with only methane flux showing significant spearman ranked correlation to methanotroph abundance, and only for the full dataset. If there was a limiting factor for methane oxidation in these cover soils, our work did not identify it. The lack of general correlation with geochemical conditions indicates that the capacity for methane oxidation in high-flux conditions is likely largely controlled by the maximum rate of methane oxidation of the resident methanotrophs. From this, the methanotrophs present and their capacity for rapid response to sudden increases in methane during hotspot formation are the key variables to optimize within bio-engineering efforts on cover soils.

There remains work to be done to determine environmental conditions supporting optimal methanotrophy at sites with high methane emissions. This work also highlights a potential future issue, where current landfill cover soil microbial communities are optimized for temperate conditions, but cover soils are already experiencing periodic windows above these communities’ maximum optimal temperatures, especially at hotspots which experience higher soil temperatures. The impact of community turnover to thermophilic lineages during hotspot events is unknown but would almost certainly result in disrupted or lowered methane oxidation rates. It is therefore important to explore the impact of increased soil temperatures due to climate change on the expected methane mitigation from landfill cover soil microbial communities.

## Supporting information

Supplemental file 1 - Spearman tests

## Acknowledgments

We are grateful to Dr. Richard Elgood from the Department of Earth and Environmental Sciences for guidance with soil gas sampling and for processing methane samples. We thank Marianne Vandergriendt from the Ecohydrology Research Group at UWaterloo for conducting soil chemistry measurements. This work was supported by an Environment and Climate Change Canada Climate Action and Awareness Fund grant EDF-CA-2012i012 to LAH and MS. LAH and MS were supported by the Canada Research Chairs. ER was supported by an NSERC USRA fellowship.

## Competing Interests

The authors state that they have no competing interests.

## Data Availability

The 16S rRNA amplicon sequence data was submitted to the NCBI SRA database under BioProject PRJNA1311329.

## Notes

### Competing Interest Statement

The authors have declared no competing interest.

