## Supplemental figures and tables for "Methanotroph Dynamics at Landfill Cover Soil Methane Emission Hotspots"

**Supplemental materials**

| | |
|---|---|
| Figures S1-S7 | p. 2-8 |
| Tables S1-S2 | p. 9-10 |

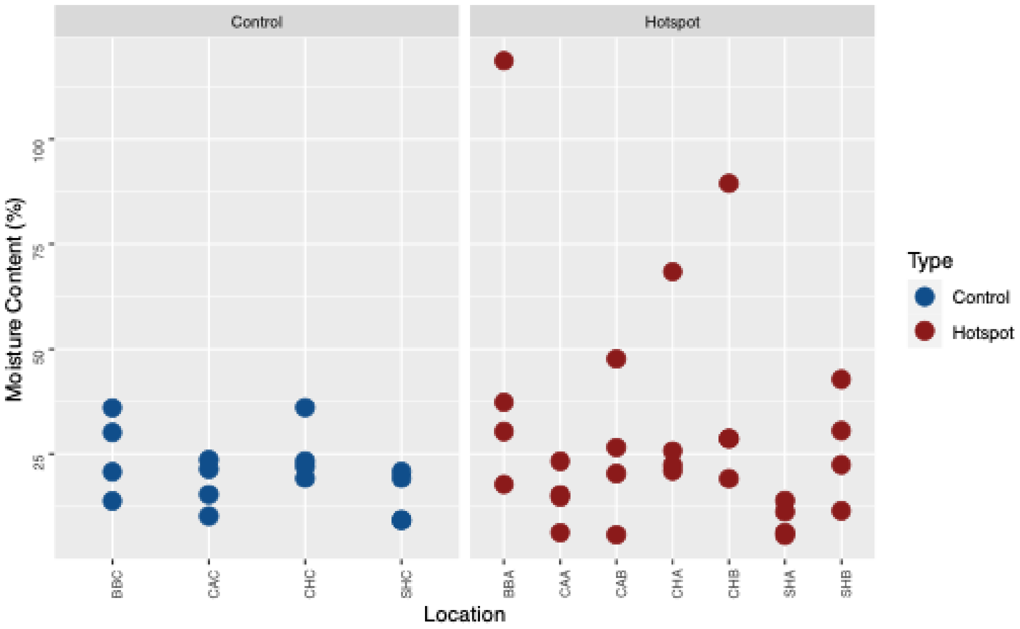


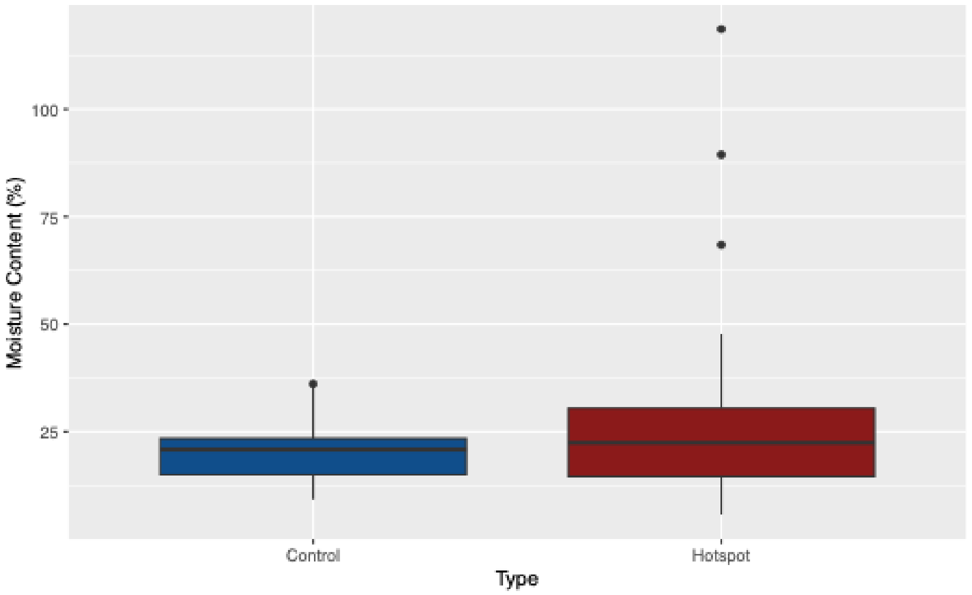


**Figure S1.** The calculated soil moisture content values (%) observed across sites as a scatter plot (A) and a Control and Hotspot site comparison as a box plot (B). Soil moisture content measurements were taken using the soil sample from the 0-meter mark along each transect, for a total of 44 samples (N=44). The Kruskal-Wallis rank sum test displayed significant ($p = 0.04785$) and non-significant ($p = 0.4792$) differences between all site locations and site type, respectively. Box plot borders indicate the first and third quartile range of values, with the line indicating the mean. Outlier values are displayed as points. Site names are as in Figure 1.

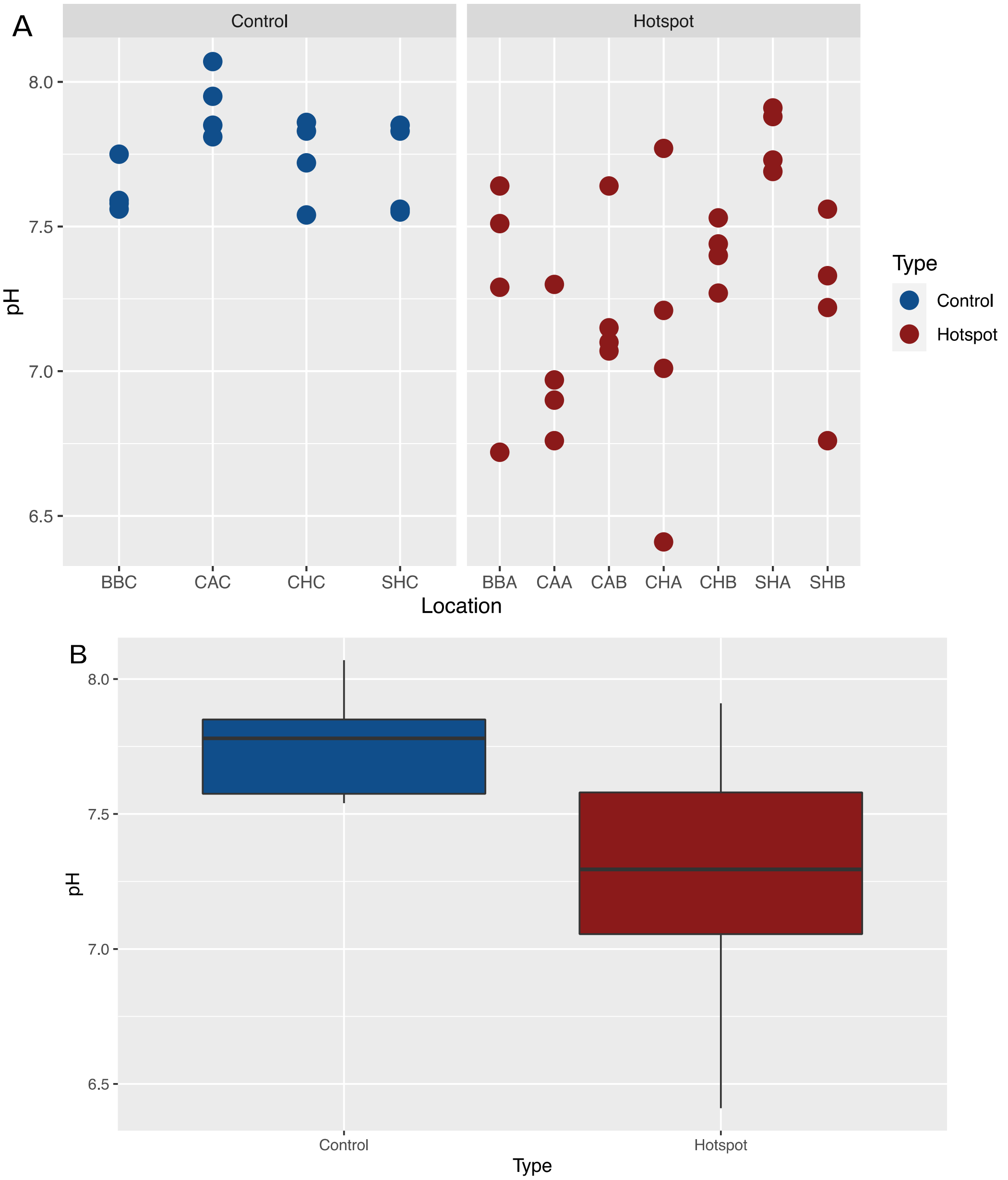


**Figure S2.** The measured pH values observed across sites as a scatter plot (A) and a Control and Hotspot comparison as a box plot (B). This measurement was taken using the soil sample of the 0-meter mark along each transect, for a total of 44 samples (N=44). Site names are as in Figure 1. The Kruskal-Wallis rank sum test depicts significant differences between all locations and between the type of site, with p values of 0.001048 and $9.434 \times 10^{-5}$, respectively. Box plot borders indicate the first and third quartile range of values, with the line indicating the mean. Outlier values are displayed as points. Site names are as in Figure 1.

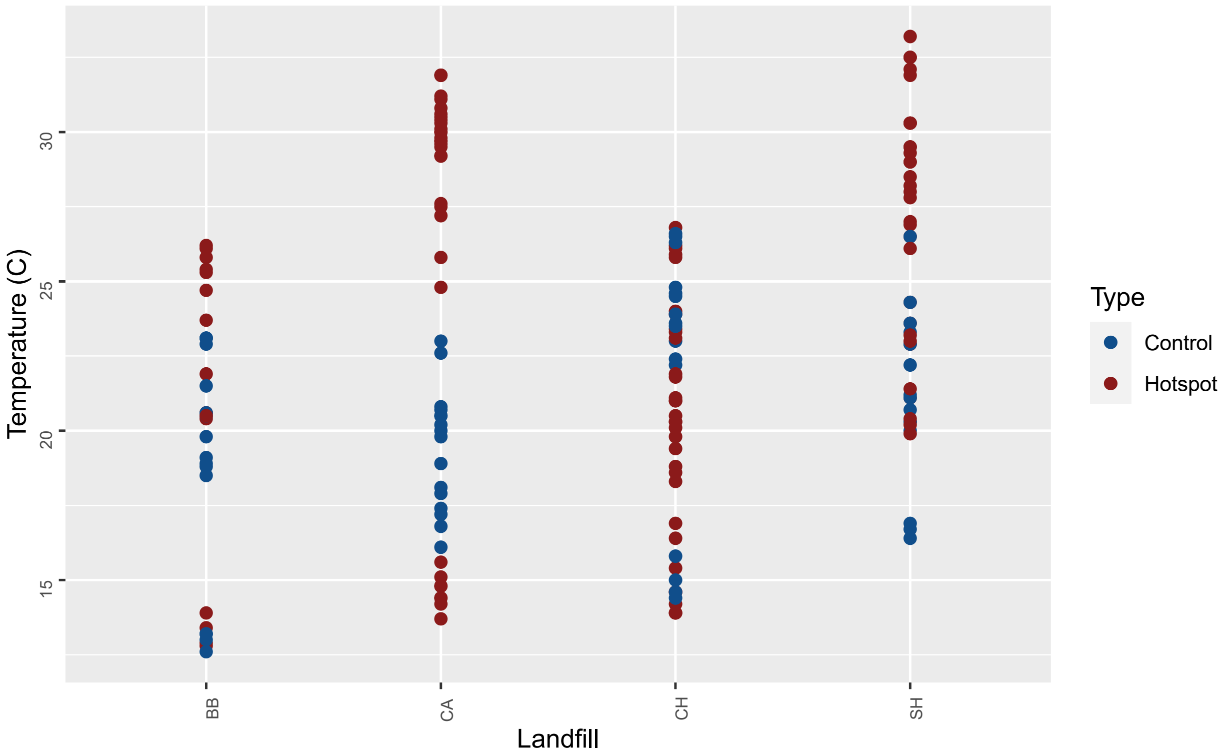


**Figure S3.** Scatter plot of the soil temperature, in degrees Celsius (C), across landfill, coloured by site type (hotspot or control) including all points along each transect for all four months (N=176). Using the Kruskal-Wallis rank sum test, there is a significant difference between site type, hotspot and control ($p = 2.11 x 10^{-5}$).

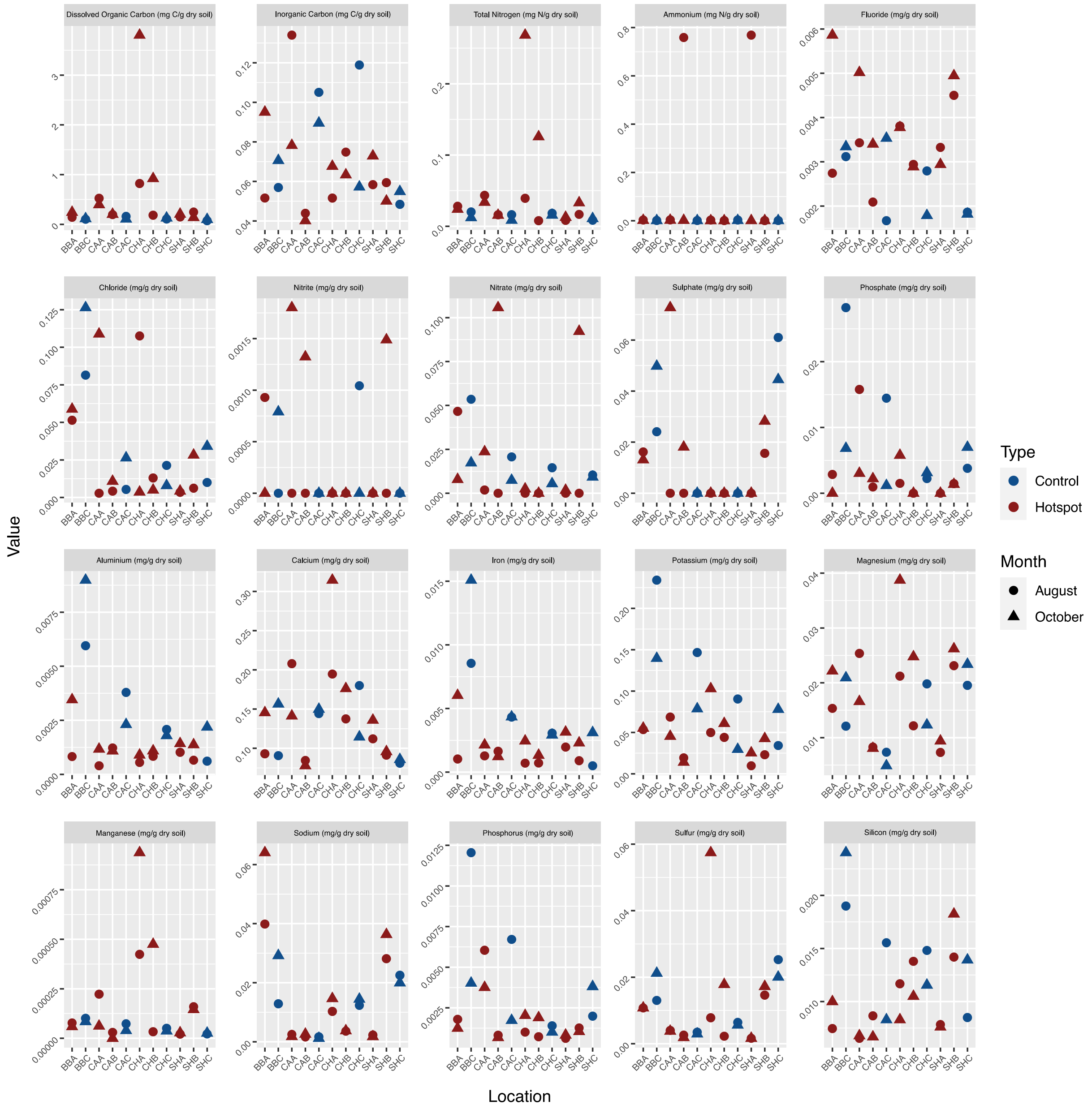


**Figure S4.** Scatter plots for the different soil chemistry measurements analysed across all sites, coloured by the sample type (hotspot or control) and differentiated by month based on the shape (N=22 for each soil chemistry parameter measured). DOC = Dissolved organic carbon. Site names are as in Figure 1.

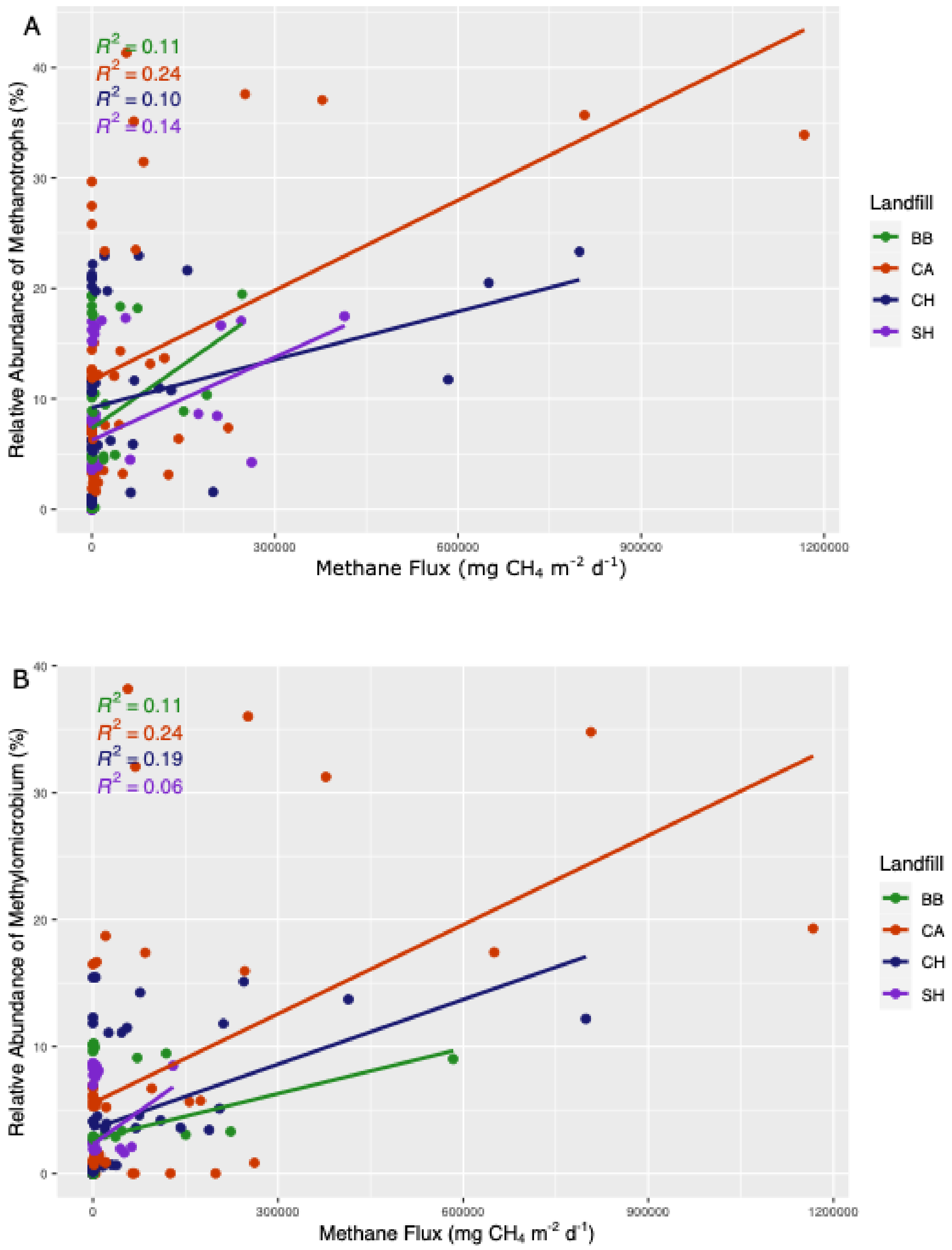


**Figure S5.** Scatter plot between the methane flux observed at each site and (A) the relative proportion of total methanotrophs or (B) the relative abundance of the dominant genus of methanotrophs observed, *Methylomicrobium*, (N=174). The calculated $R^2$ values represent the strength of the correlation, visualized as linear trendlines for each landfill.

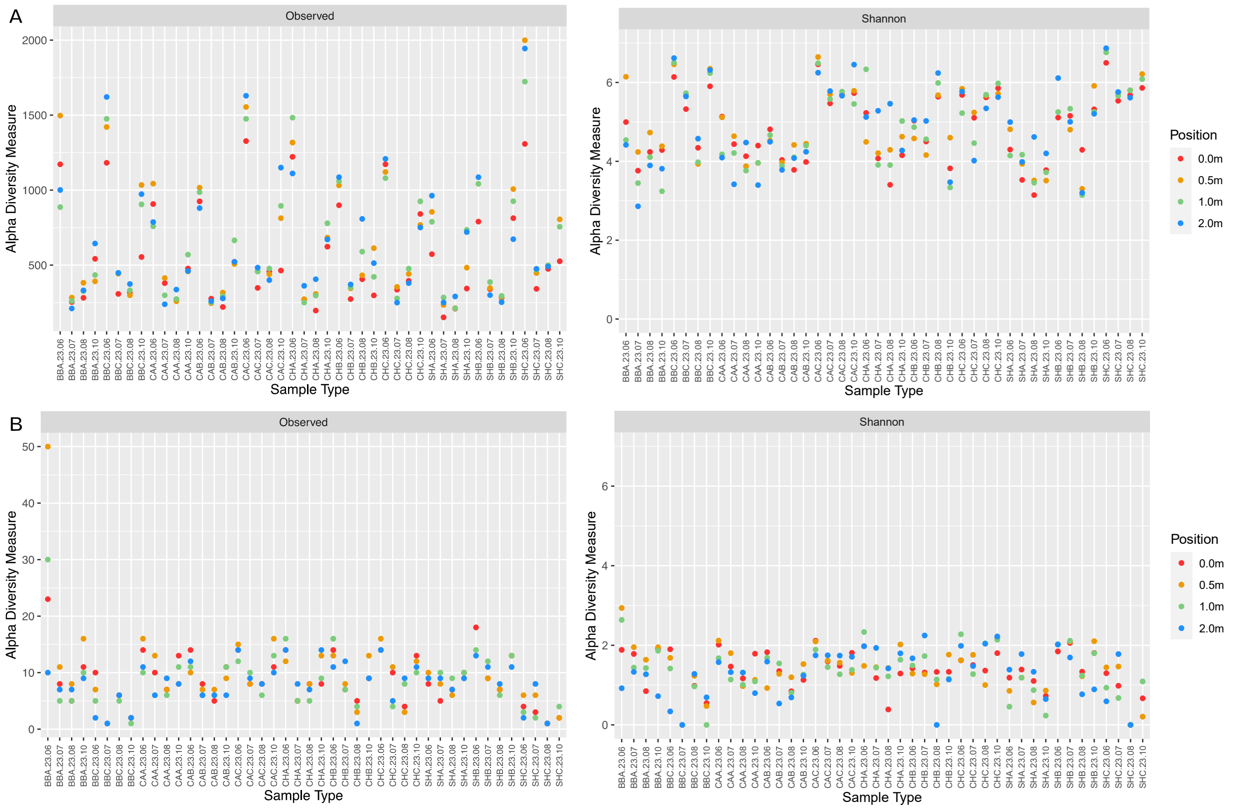


**Figure S6.** The Observed (left) and Shannon (right) index for the total community diversity (A) and methanotroph diversity (B) for each sample, note different axes for the observed or richness of the community. Samples each have four points along the sampling transect (0 m, 0.5 m, 1 m, and 2 m), coloured according to distance from the hotspot or control location. Sample names describe the landfill (BB, CA, CH, and SH), the site (A, B, and C), the year and month of sampling (23.06, 23.07, 23.08, and 23.10), as well as the distance along the transect in meters (0.0, 0.5, 1.0, and 2.0).

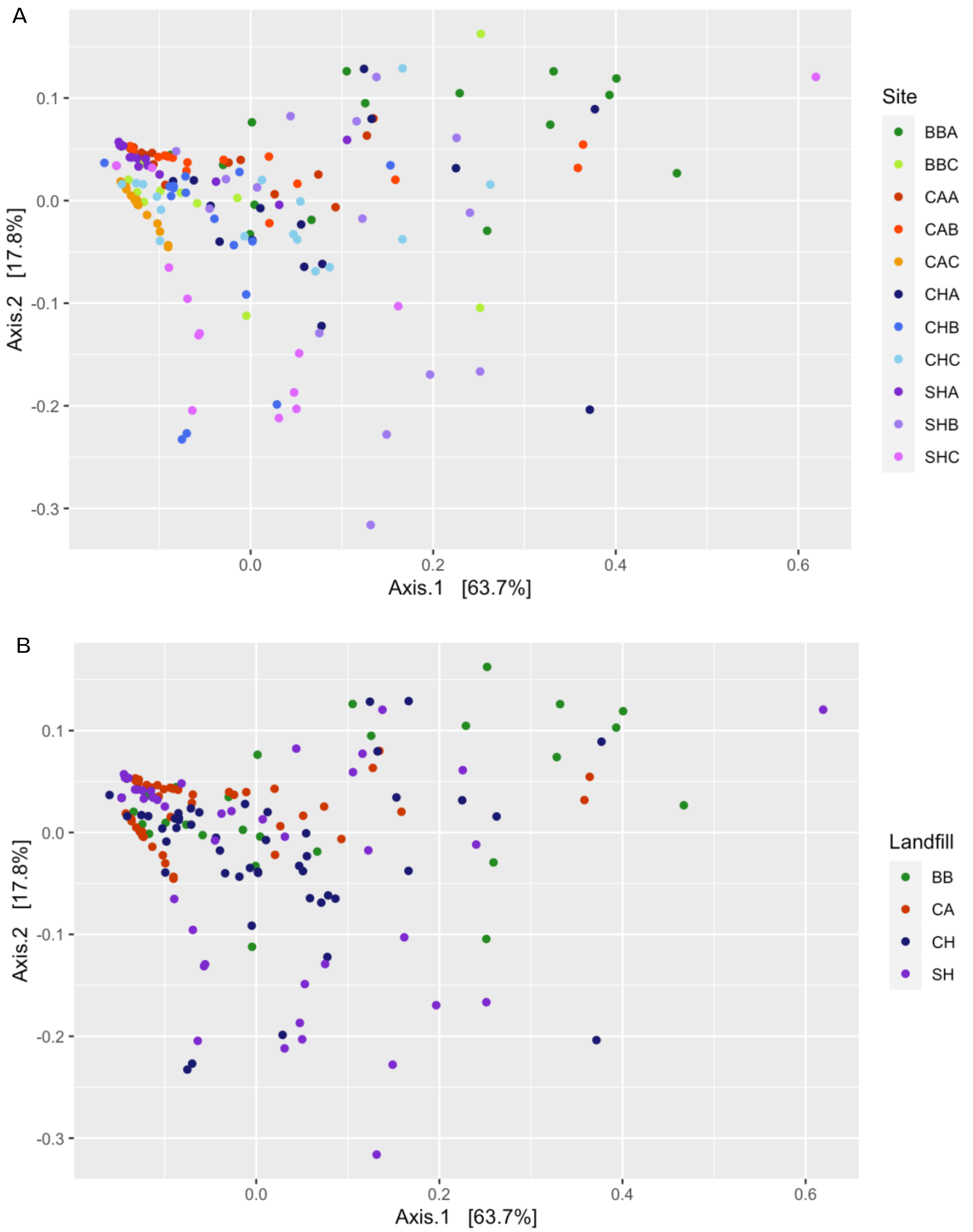


**Figure S7.** PCoA plot of the weighted Unifrac diversity metric for methanotroph-only data across all samples. Colours distinguish between the different sampling sites (A) and the four landfills (B) (N=174). Site names are as in Figure 1.

**Table S1**: Spearman correlation coefficients (rho) and Benjamini-Hochberg-corrected [p-values] for significantly correlated pairs of variables. For the full set of pairwise comparisons, see Supplementary Data File 1. A p-value below 0.05 was considered significant. For the full dataset (F), n=22-174 depending on the variables. For hotspots only (H), n=14-111. For control sites (C), n=8-63.

| Variable Pair | Dataset | Rho (P-value) | | |
|---|---|---|---|---|
| | | **ALL** | **HOTSPOT** | **CONTROL** |
| Aluminum and Iron | F, H, C | 0.93 (9.82E-08) | 0.78 (0.02) | 0.98 (0.011) |
| Flux and Soil gas | F, H | 0.73 (7.15E-14) | 0.55 (1.34E-05) | |
| Flux and Soil temperature | F, H | 0.41 (2.25E-06) | 0.33 (0.015) | |
| Manganese and TN | F, H | 0.81 (0.00031) | 0.88 (0.0025) | |
| Phosphorus and Potassium | F, H | 0.81 (0.00031) | 0.8 (0.017) | |
| Flux and Nitrate | F, H | -0.77 (0.001) | -0.74 (0.034) | |
| Magnesium and Sulfur | F, H | 0.75 (0.0018) | 0.87 (0.003) | |
| Sodium and Sulfur | F, H | 0.75 (0.0018) | 0.72 (0.047) | |
| Nitrate and Nitrite | F, H | 0.69 (0.01) | 0.79 (0.019) | |
| Nitrate and Sulphate | F, H | 0.65 (0.025) | 0.75 (0.032) | |
| Magnesium and Manganese | F, H | 0.64 (0.028) | 0.88 (0.0025) | |
| Magnesium and TN | F, H | 0.63 (0.033) | 0.81 (0.016) | |
| Soil gas and Soil temperature | Full | 0.47 (2.25E-06) | | |
| Phosphate and Phosphorus | Full | 0.79 (0.0006) | | |
| Calcium and DIC | Full | 0.7 (0.0069) | | |
| Flux and Methanotroph | Full | 0.28 (0.0069) | | |
| Calcium and Manganese | Full | 0.61 (0.039) | | |
| Chloride and Sulphate | Full | 0.62 (0.039) | | |
| DOC and TN | Full | 0.62 (0.039) | | |
| Magnesium and Sodium | Full | 0.62 (0.039) | | |
| Nitrate and Soil gas | Full | -0.6 (0.044) | | |
| Nitrite and Sulphate | Hotspots | | 0.88 (0.0025) | |
| Manganese and Sulfur | Hotspots | | 0.85 (0.0074) | |
| Phosphorus and TN | Hotspots | | 0.83 (0.012) | |
| Calcium and Potassium | Hotspots | | 0.82 (0.013) | |
| Potassium and TN | Hotspots | | 0.8 (0.017) | |
| Flux and Nitrite | Hotspots | | -0.78 (0.02) | |
| Manganese and Potassium | Hotspots | | 0.78 (0.02) | |
| Flux and Sulphate | Hotspots | | -0.78 (0.021) | |
| Magnesium and Phosphorus | Hotspots | | 0.76 (0.026) | |
| Sulfur and TN | Hotspots | | 0.76 (0.026) | |
| Manganese and Phosphorus | Hotspots | | 0.75 (0.03) | |
| Magnesium and Potassium | Hotspots | | 0.75 (0.031) | |
| Calcium and DOC | Hotspots | | 0.72 (0.049) | |

**Table S2.** Results from statistical tests between site (XXA, XXB, and XXC), landfill (SH, CH, CA, and BB), and site type (Hotspot vs Control) for the total community microbial diversity metrics (top) and the methanotroph fraction (bottom). Significance was assessed using an p value of 0.05. Statistical tests include multivariate permutational analysis of variance (PERMANOVA), multivariate permutational analysis of dispersion (PERMDISP), and Kruskal-Wallis test. Matrices included unweighted and weighted Unifrac, Bray-Curtis, Shannon index, and observed features. Significant tests are highlighted in grey.

| Metric | Site | | | Landfill | | | Site type | | |
|---|---|---|---|---|---|---|---|---|---|
| | PERM-ANOVA | PERM-DISP | Kruskal-Wallis | PERM-ANOVA | PERM-DISP | Kruskal-Wallis | PERM-ANOVA | PERM-DISP | Kruskal-Wallis |
| **Total community** | | | | | | | | | |
| **Unweighted** | 0.001 | 0.001 | | 0.001 | 0.018 | | 0.001 | 0.001 | |
| **Weighted** | 0.001 | 0.001 | | 0.001 | 0.001 | | 0.001 | 0.001 | |
| **Bray-Curtis** | 0.002 | 0.741 | | 0.001 | 0.739 | | 0.03 | 0.557 | |
| **Shannon** | | | $5.03 \times 10^{-16}$ | | | 0.784 | | | $7.76 \times 10^{-19}$ |
| **Observed** | | | 0.002 | | | 0.885 | | | $4.04 \times 10^{-6}$ |
| **Methanotrophs** | | | | | | | | | |
| **Unweighted** | 0.001 | 0.254 | | 0.058 | 0.639 | | 0.001 | 0.3 | |
| **Weighted** | 0.016 | 0.013 | | 0.021 | 0.02 | | 0.058 | 0.089 | |
| **Bray-Curtis** | 1.0 | 0.236 | | 0.107 | 0.749 | | 0.57 | 0.941 | |
| **Shannon** | | | 0.059 | | | 0.132 | | | 0.269 |
| **Observed** | | | 0.024 | | | 0.025 | | | 0.142 |
